# A flexible network of Vimentin intermediate filaments promotes the migration of amoeboid cancer cells through confined environments

**DOI:** 10.1101/788810

**Authors:** Sara M. Tudor, Sandrine B. Lavenus, Jeremy S. Logue

## Abstract

The spread of tumor cells to distant sites is promoted by their ability to switch between mesenchymal and amoeboid (bleb-based) migration. Because of this, inhibitors of metastasis must account for each motility mode. To this end, here we determine the precise role of the Vimentin intermediate filament system in regulating the migration of amoeboid human cancer cells. Vimentin is a classic marker of epithelial to mesenchymal transition and is therefore, an ideal target for a metastasis inhibitor. However, the role of Vimentin in amoeboid migration has not been determined. Since amoeboid, leader bleb-based migration occurs in confined spaces and Vimentin is known to be a major determinant of cell mechanical properties, we hypothesized that a flexible Vimentin network is required for fast amoeboid migration. This was tested using our PDMS slab-based approach for the confinement of cells, RNAi, over-expression, pharmacological treatments, and measurements of cell stiffness. In contrast to Vimentin RNAi, inducing the bundling of Vimentin was found to inhibit fast amoeboid migration and proliferation. Importantly, these effects were independent of changes in actomyosin contractility. Collectively, our data supports a model whereby the perturbation of cell mechanical properties by Vimentin bundling inhibits the invasive properties of cancer cells.

## Introduction

Cell migration is required for embryonic development, immune surveillance, and wound healing in healthy individuals. However, the uncontrolled migration of tumor cells to distant sites is a hallmark of metastasis and is associated with poor prognosis. In recent years, it has been demonstrated that cancer cells can switch between focal adhesion (mesenchymal) and bleb-based (amoeboid) migration modes. This is important because blocking metastasis will require that each mode of migration be targeted. To this aim, here we determine the role of a well-established regulator of mesenchymal migration, the Vimentin Intermediate Filament (VIF) cytoskeleton, in regulating the amoeboid migration of cancer cells.

The switch from a predominantly Keratin to Vimentin expression pattern is a classic marker of Epithelial to Mesenchymal Transition (EMT). Accordingly, Vimentin is known to increase the size and strength of focal adhesions (1). In contrast, the role of Vimentin in amoeboid (blebbing) cells has not been determined. Using *in vivo* and *in vitro* approaches, it has been shown that highly contractile (metastatic) cancer cells will switch from a mesenchymal to “fast amoeboid” mode of migration in response to physically confining environments, such as those found in micro-lymphatics/capillaries and perivascular spaces (2-6). Additionally, certain drug treatments including, Matrix Metallo**<u>p</u>**rotease (MMP) and tyrosine kinase inhibitors (e.g., Dasatinib), will induce a switch to bleb-based migration (7-9). Fast amoeboid migration relies on the formation of what we termed a leader bleb (2). In confined environments, leader blebs are typically very large and stable blebs containing a rapid cortical actomyosin flow (2-5). Whereas mesenchymal cells utilize integrin-Extra**<u>c</u>**ellular Matrix (ECM) interactions for migration, fast amoeboid or Leader Bleb-Based Migration (LBBM) only requires friction between the cortical actomyosin flow and the extracellular environment (5). This property likely promotes the invasive properties of cancer cells *in vivo*.

Because metastasis requires that cells migrate within the confines of tissues, we hypothesized that confined cancer cell migration (i.e., LBBM) requires a flexible intermediate filament network. Unlike Keratin, which stiffens by bundling in response to force (i.e., strain stiffens), Vimentin remains unbundled and flexible (10). Moreover, photobleaching experiments have shown that Vimentin undergoes subunit exchange an order of magnitude faster than Keratin and are therefore, considered to be more dynamic (11). Recently, a statin used for lowering blood cholesterol, Simvastatin, was identified in a screen for Vimentin binding molecules (12). In contrast to other statins, such as Pravastatin, Simvastatin was found to bind the sides of Vimentin and induce bundling. In cell-based assays, Simvastatin was shown to block the proliferation of adrenal carcinoma cells, possibly because Vimentin bundling inhibits its degradation required for cell division (13). Importantly, Simvastatin binds Vimentin with high specificity, as opposed to other molecules (e.g., Withaferin A) that effect other components of the cytoskeleton (14,15). Here, by combining Simvastatin with our recently described approach for the confinement of cells, we describe the precise role of a flexible (unbundled) Vimentin network in amoeboid human cancer cells (16).

Our data show that the concentration of Vimentin and its bundling are potent regulators of mesenchymal and amoeboid migration, mechanics, and the survival of human cancer cells in confinement. Collectively, this work sheds new light on the potential of Vimentin as a therapeutic target.

## Results

Because a high level of Vimentin expression is correlated with hematogenous metastasis within a wide array of melanoma samples, we set out to determine the localization of Vimentin in melanoma A375-M2 cells (17). Moreover, this highly metastatic sub-line has been observed by intravital imaging to undergo amoeboid migration in tumors (18). Using Vimentin tagged on its C-terminus with FusionRed, Vimentin-FusionRed, and transient transfection we determined the localization of Vimentin in A375-M2 cells by live high-resolution microscopy (Fig. 1). In cells adhered to fibronectin coated glass, an isotropic network of Vimentin was concentrated near the cell center (Fig. 1A, *left*). Similarly, in non-adherent (blebbing) cells on uncoated glass, Vimentin surrounded the nucleus and was excluded from blebs (Fig. 1A, *middle*). In order to evaluate the localization of Vimentin in cells with leader blebs, we promoted the conversion of A375-M2 cells to this morphology by confinement using our Poly**<u>d</u>**i**<u>m</u>**ethyl**<u>s</u>**iloxane (PDMS) slab-based approach (16). This involves placing cells between a Bovine Serum Albumin (BSA; 1%) coated slab of PDMS and cover glass, which is held at a defined height by beads with a diameter of ∼3 µm. This confinement height was shown to be optimal for stimulating the transition to fast amoeboid migration (3). Using this approach, we found that Vimentin was kept entirely within the cell body as opposed to leader blebs (Fig. 1A, *right* & Movie S1). This is significant because the cell body is the principle source of resistance to the cortical actomyosin flow in leader blebs, limiting LBBM speed (3).

**Figure 1.**
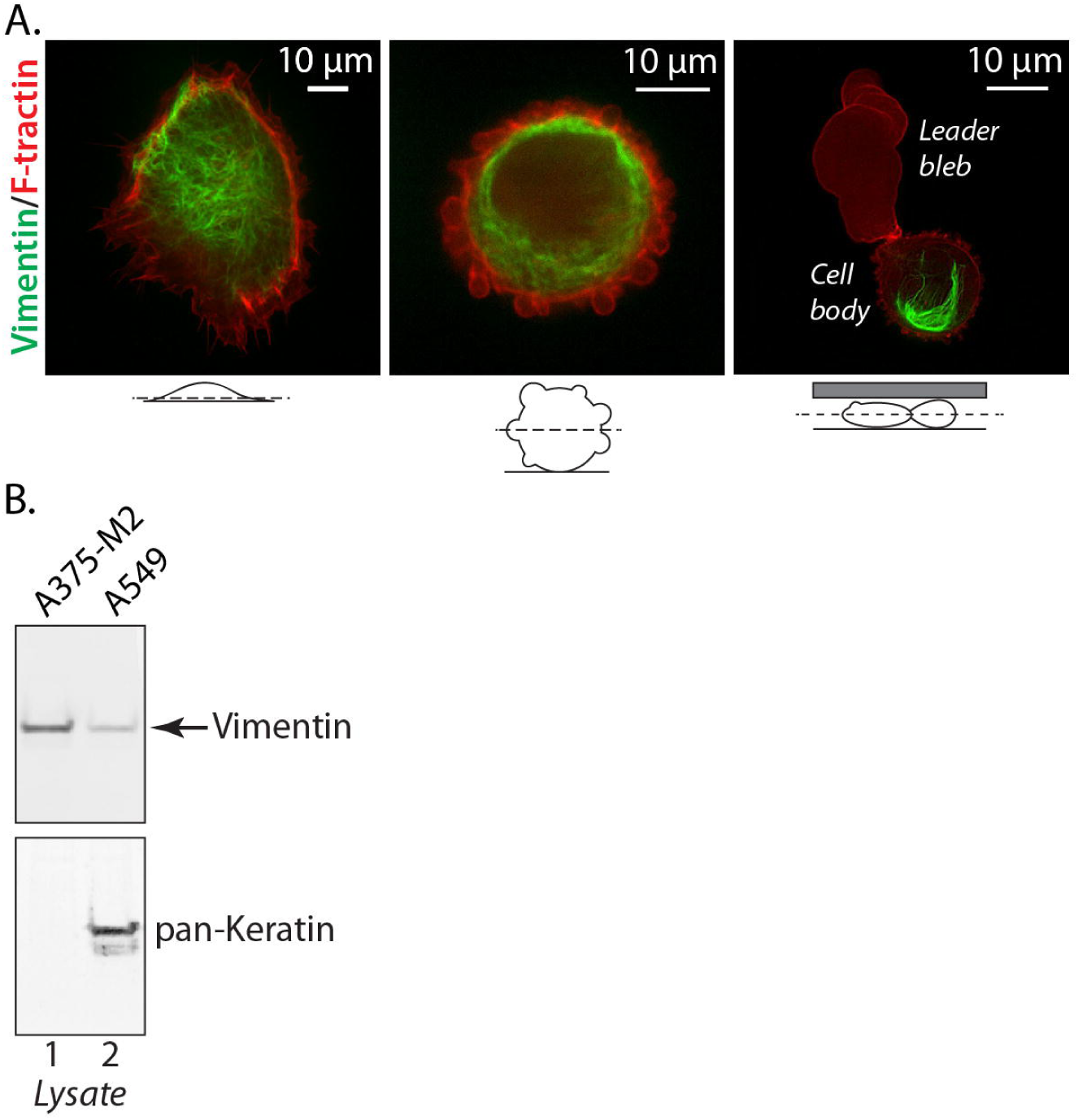
Vimentin localizes to the cell body of leader bleb forming cells. **A.** A375-M2 cells adhered to fibronectin (left), uncoated glass (blebbing; middle), and confined under PDMS (forming a leader bleb; right) transiently expressing Vimentin-FusionRed. **B.** Lysates from A375-M2 and A549 cells probed for endogenous Vimentin and pan-Keratin. All data are representative of at least three independent experiments. See also supplemental movie 1.

To directly test the notion that Vimentin concentration in the cell body limits LBBM speed, we depleted A375-M2 cells of Vimentin using a Locked Nucleic Acid (LNA), which offer enhanced specificity and stability over traditional small interfering RNAs (siRNAs) (19). Because of the long half-life of Vimentin, cells were incubated with LNAs for 5 days to achieve a ∼90% reduction in protein levels (Fig. 2B). Moreover, because these cells predominantly express Vimentin, they are an ideal (simplified) model for defining the role of intermediate filaments in LBBM (Fig. 1B). Using our PDMS slab-based approach, LBBM was quantitatively evaluated for LNA treated cells by live imaging over 5 hr. Strikingly, in cells depleted of Vimentin, the speed of LBBM was increased over control by ∼50% (Fig. 2E-F & Movie S2), whereas directionality remained the same for each group (Fig. S1A). Quantitation proved that leader bleb area, which is defined as the single largest bleb within a given frame, is close to double the size of control (Fig. 2C). Interestingly, quantitation of cell body area found that in Vimentin RNAi cells, the cell body area was decreased by over 25% (Fig. 2D). This result is consistent with the location of Vimentin in these cells, which may limit the degree to which cortical actomyosin is able to contract the cell body. Consequently, more cytoplasm from the cell body can enter leader blebs, increasing their size. Strikingly, the nucleus in Vimentin RNAi cells was observed to undergo large shape changes, which may reflect an increase in the degree of force transmitted to the nucleus from cortical actomyosin (Fig. 2A). A more than 25% increase in the number of Vimentin RNAi cells undergoing apoptosis is consistent with reports of nuclear rupture and DNA damage in confined cells (Fig. S2A) (20,21). To test the hypothesis that Vimentin regulates the stiffness of A375-M2 cells, we used an approach described by the Piel Lab (Institut Curie) that involves compressing cells between two Poly**<u>a</u>**crylamide (PA; 1 kPa) gels (Fig. 2G) (3). Using this approach, cell height divided by the diameter (h/d), which is a function of the opposing force, is used to define the “cell stiffness.” Consistent with other reports, we find that cells depleted of Vimentin are ∼25% softer than control cells (Fig. 2H). Because cortical actomyosin is also expected to regulate stiffness, we confirmed that the level of active (phosphorylated) Regulatory Light Chain (p-RLC) is not affected by Vimentin RNAi (Fig. 2I). Therefore, Vimentin expression increases cell stiffness to limit migration in confined environments.

**Figure 2.**
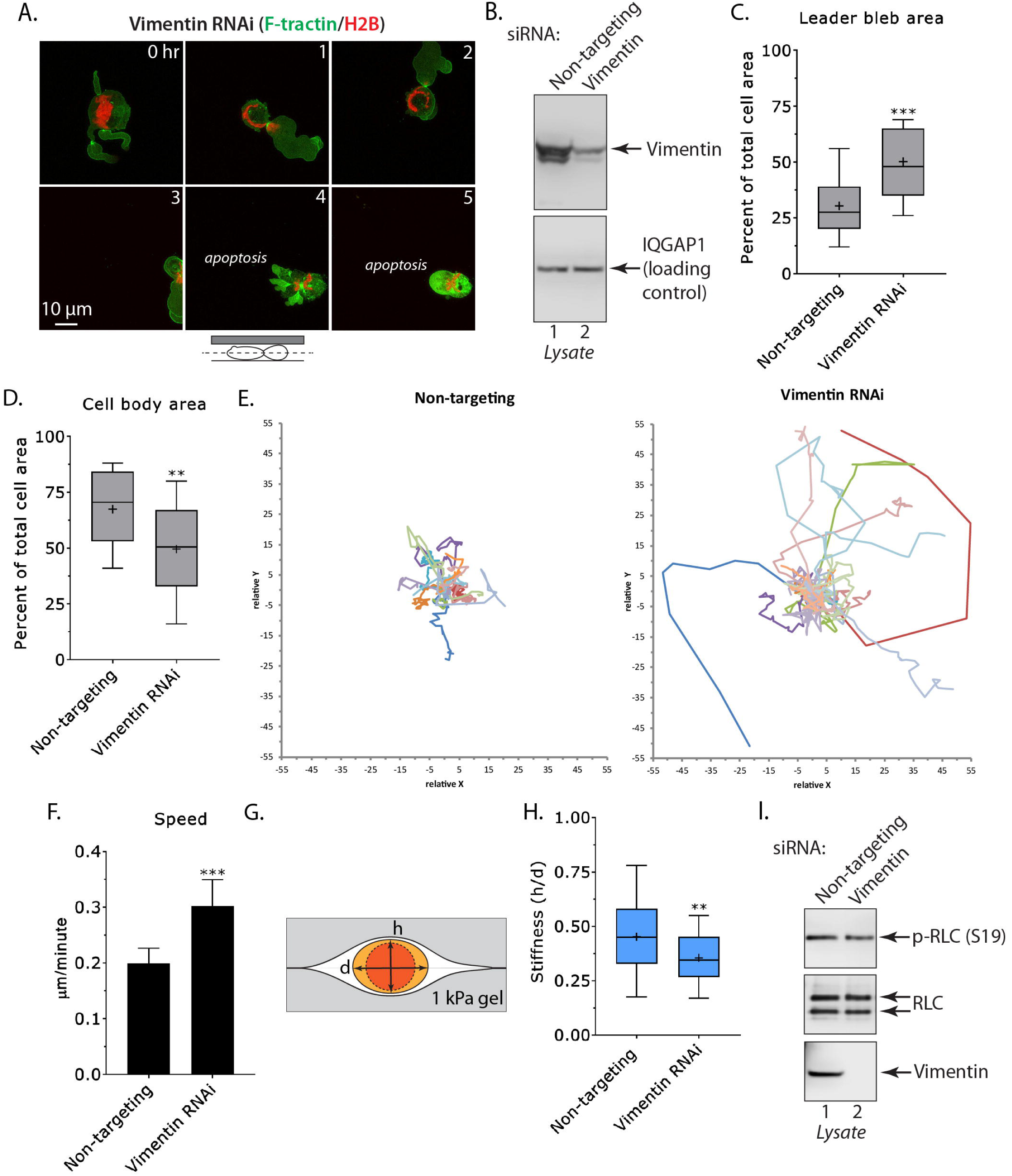
RNAi of Vimentin promotes rapid leader bleb-based migration. **A.** Montage of an A375-M2 cell under PDMS depleted of Vimentin by RNAi expressing EGFP-F-tractin and H2B-FusionRed. **B.** Western blot confirming the depletion of Vimentin by RNAi in A375-M2 cells. **C-D.** Quantitative evaluation of leader bleb (C) and cell body (D) area in control (non-targeting) and Vimentin RNAi cells. Statistical significance was determined by two-tailed (unpaired) Student’s t-tests. **E.** Migration tracks for non-targeting (left; n=40) and Vimentin RNAi (right; n=43) cells under PDMS. **F.** Instantaneous speeds for non-targeting and Vimentin RNAi cells. Statistical significance was determined by an F test. Error is SEM. **G.** Cartoon of the gel sandwich approach for measuring cell stiffness. **H.** Cell stiffness for non-targeting (n=77) and Vimentin RNAi (n=30) cells. Statistical significance was determined by a two-tailed (unpaired) Student’s t-test. **I.** Western blots of endogenous phosphorylated Regulatory Light Chain (p-RLC; S19) and RLC in non-targeting and Vimentin RNAi cells. Tukey box plots in which “+” and line denote the mean and median, respectively. All data are representative of at least three independent experiments. See also supplemental figure 1, 2A, and movie 2. * - p ≤ 0.05, ** - p ≤ 0.01, and *** - p ≤ 0.001

Because depleting A375-M2 cells of Vimentin had a striking effect on confined migration, we set out to quantitatively evaluate the effect of Vimentin Over-Expression (OE) on LBBM. To this end, we transiently transfected cells with Vimentin-FusionRed and performed live imaging (Fig. 3A). In contrast to Vimentin RNAi, increasing the concentration of Vimentin did not significantly affect leader bleb or cell body area (Fig. 3B-C). However, the cell body area trended towards being larger in Vimentin OE cells, which is consistent with the concept that Vimentin limits the compressibility of the cell body (Fig. 3C). Additionally, LBBM speed trended towards being lower in Vimentin OE cells when compared to the expression of EGFP alone (Fig. 3D). Directionality over time was unchanged in Vimentin OE cells, whereas we observed a slight increase in the number of apoptotic cells (Fig. S1B & S2B). Interestingly, using the gel sandwich approach, cell stiffness was increased by ∼50% after Vimentin OE (Fig. 3E). Therefore, increasing the expression of Vimentin can influence the mechanics of A375-M2 cells. The absence of larger effects on LBBM may be a consequence of the high endogenous level of Vimentin in A375-M2 cells (Fig. 1B & https://www.proteinatlas.org/); therefore, an incremental increase in Vimentin levels may not be sufficient to affect LBBM.

**Figure 3.**
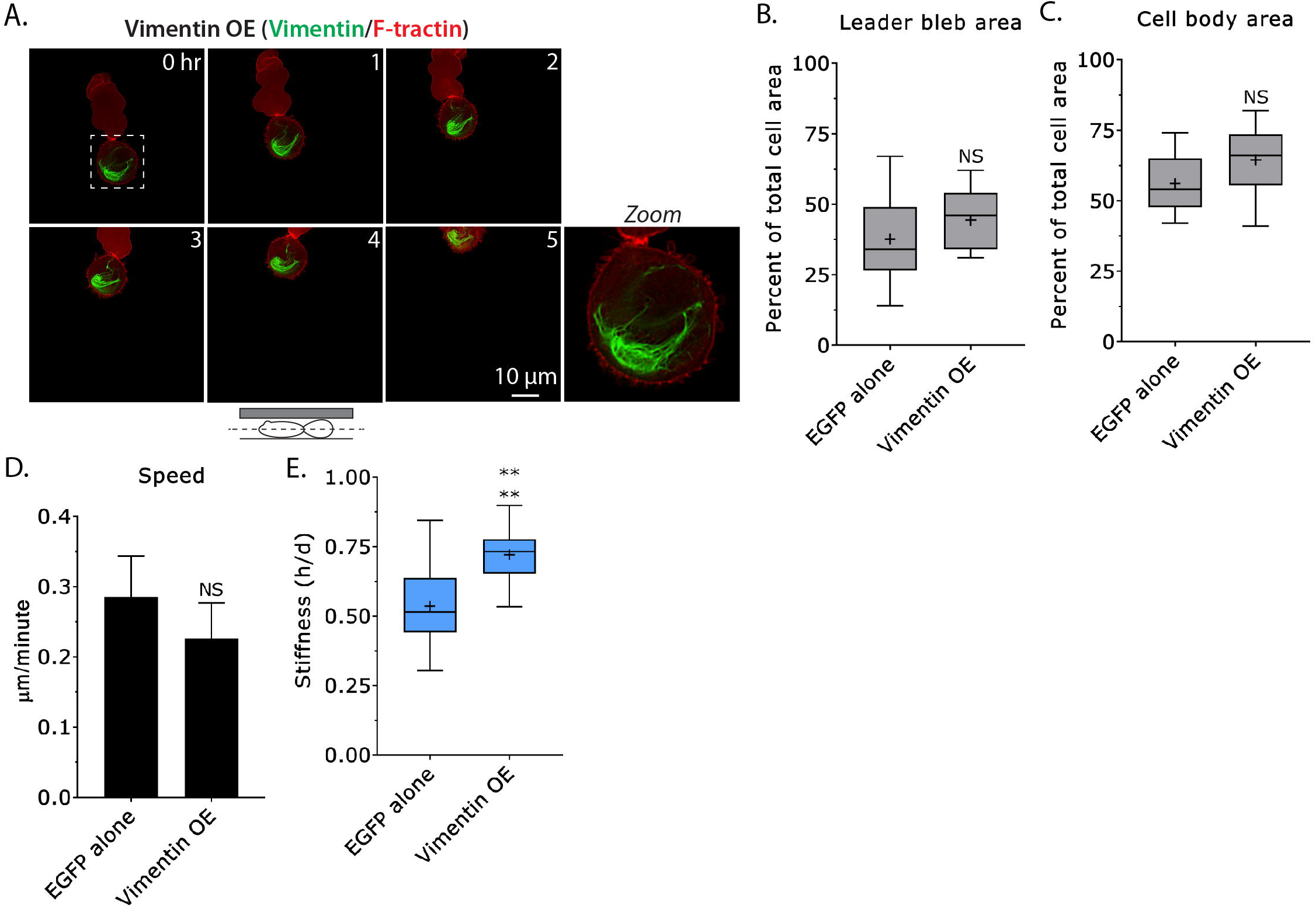
Over-expressing Vimentin in confined cells. **A.** Montage of an A375-M2 cell over-expressing Vimentin and the marker of Filamentous actin (F-actin), F-tractin, under PDMS. **B-C.** Quantitative evaluation of leader bleb (B) and cell body (C) area for cells over-expressing EGFP alone and Vimentin-FusionRed. Statistical significance was determined by two-tailed (unpaired) Student’s t-tests. **D.** Instantaneous speeds for cells over-expressing EGFP alone (n=41) and Vimentin-FusionRed (n=42). Statistical significance was determined by an F test. Error is SEM. **E.** Cell stiffness for EGFP alone (n=34) and Vimentin-FusionRed (n=31) over-expressing cells. Statistical significance was determined by a two-tailed (unpaired) Student’s t-test. Tukey box plots in which “+” and line denote the mean and median, respectively. All data are representative of at least three independent experiments. See also supplemental figure 2B, and movie 1. * - p ≤ 0.05, ** - p ≤ 0.01, and *** - p ≤ 0.001

Next, we set out to determine if modulating the architecture of Vimentin can impact LBBM. Recently, a potent and selective inducer of Vimentin bundling was identified by an image-based screen (12). In this screen, the cholesterol lowering statin, Simvastatin, was shown to directly bind the sides of Vimentin and likely through limiting electrostatic repulsion between filaments, is able to induce filament bundling (12). This result is significant because in large cohort studies, patients taking statins for cholesterol have decreased cancer associated morbidity (22). Therefore, we treated A375-M2 cells with Simvastatin (10 µM) and performed live imaging of Vimentin-FusionRed. Compared to Vehicle treated (DMSO), Vimentin was observed to progressively bundle in cells treated with Simvastatin (Fig. 4A-B & Movies 3-4). To quantitatively evaluate this affect we measured the total area of the Vimentin network in Vehicle, Simvastatin, and Pravastatin treated cells. The related statin, Pravastatin, is used here for comparison with Simvastatin. Using this approach, the average area of the Vimentin network was decreased by ∼35% in Simvastatin treated cells, whereas treating cells with Pravastatin did not have a significant effect (Fig. 4C). Next, we quantified the speed of LBBM for Vehicle, Simvastatin, and Pravastatin treated cells. This analysis showed that cells treated with Simvastatin were ∼60% slower than those treated with Vehicle and Pravastatin (Fig. 4D). To evaluate the effect of Simvastatin on cell stiffness, we again used the gel sandwich approach. For cells treated with Simvastatin, we observed a small decrease (∼10%) in cell stiffness, whereas treating cells with Pravastatin had no effect when compared to Vehicle (Fig. 4E). Because the bundling of Vimentin causes the network to collapse into a small area of the cytoplasm, this result might be expected since the gel sandwich assay measures the stiffness of the entire cell. However, since bundled intermediate filament networks are known to be much stiffer, a local (large) increase in mechanical properties is expected (10). In line with this concept, we observe leader blebs to pull against sites of Vimentin bundling in the cell body (Fig. 4B & Movie 4). Because the actomyosin cytoskeleton is also expected to effect cell stiffness, we also measured the level of active myosin (p-RLC) in drug treated cells. By Western blotting, we confirmed that treatment with Vehicle, Simvastatin, and Pravastatin did not affect the level of p-RLC (Fig. 4F). Moreover, the concentration of Vimentin was unchanged in these cells (Fig. 4F). Therefore, by inducing Vimentin bundling with Simvastatin, migration in confined environments is inhibited. As tissue culture cells obtain cholesterol from serum, statin treatment is not expected to alter the level of intracellular cholesterol, which is a critical component of the plasma membrane (23). Therefore, our data supports a model whereby local stiffening of the cell body by Vimentin bundling inhibits LBBM (Fig. 5E).

**Figure 4.**
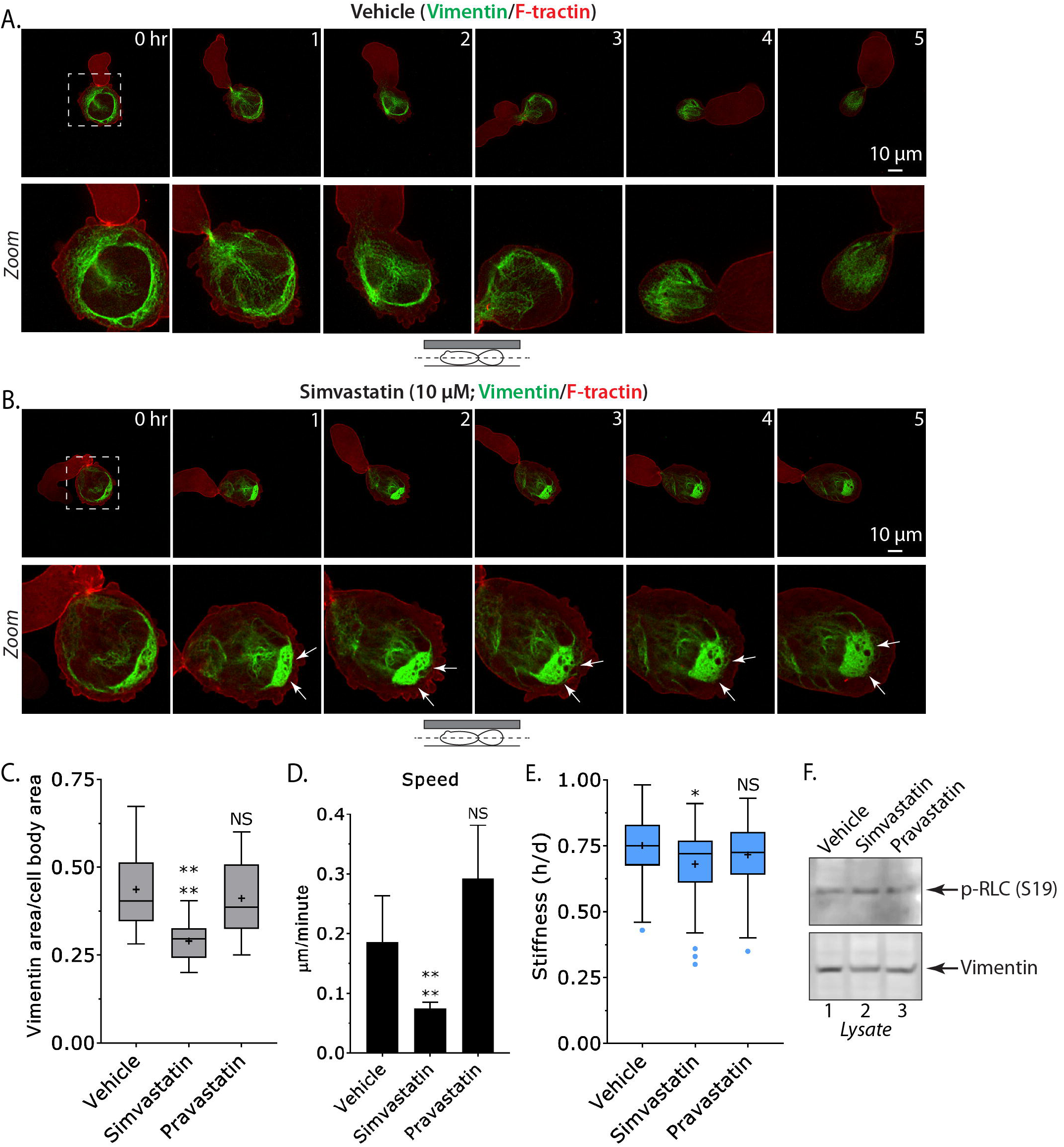
Vimentin bundling inhibits leader bleb-based migration. **A-B.** Montage of an A375-M2 cell treated with Vehicle (A; DMSO) or Simvastatin (B; 10 µM) expressing EGFP-F-tractin and Vimentin-FusionRed under PDMS. Arrows point to areas of Vimentin bundling in the cell body. **C.** Quantitative evaluation of Vimentin bundling for Vehicle (n=39), Simvastatin (n=26), and Pravastatin (n=27) treated cells. Statistical significance was determined by an ordinary one-way ANOVA followed by a post-hoc multiple comparisons test. **D.** Quantitative evaluation of instantaneous speeds for Vehicle (n=28), Simvastatin (n=19), and Pravastatin (n=20) treated cells. Statistical significance was determined by F tests. Error is SEM. **E.** Cell stiffness measurements for Vehicle (n=77), Simvastatin (n=63), and Pravastatin (n=62). Statistical significance was determined by a Kruskal-Wallis test followed by a post-hoc multiple comparisons. **F.** Western blots of endogenous p-RLC, RLC, and Vimentin in drug treated cells. Tukey box plots in which “+” and line denote the mean and median, respectively. All data are representative of at least three independent experiments. See also supplemental figure 2C, and movies 3-4. * - p ≤ 0.05, ** - p ≤ 0.01, and *** - p ≤ 0.001

**Figure 5.**
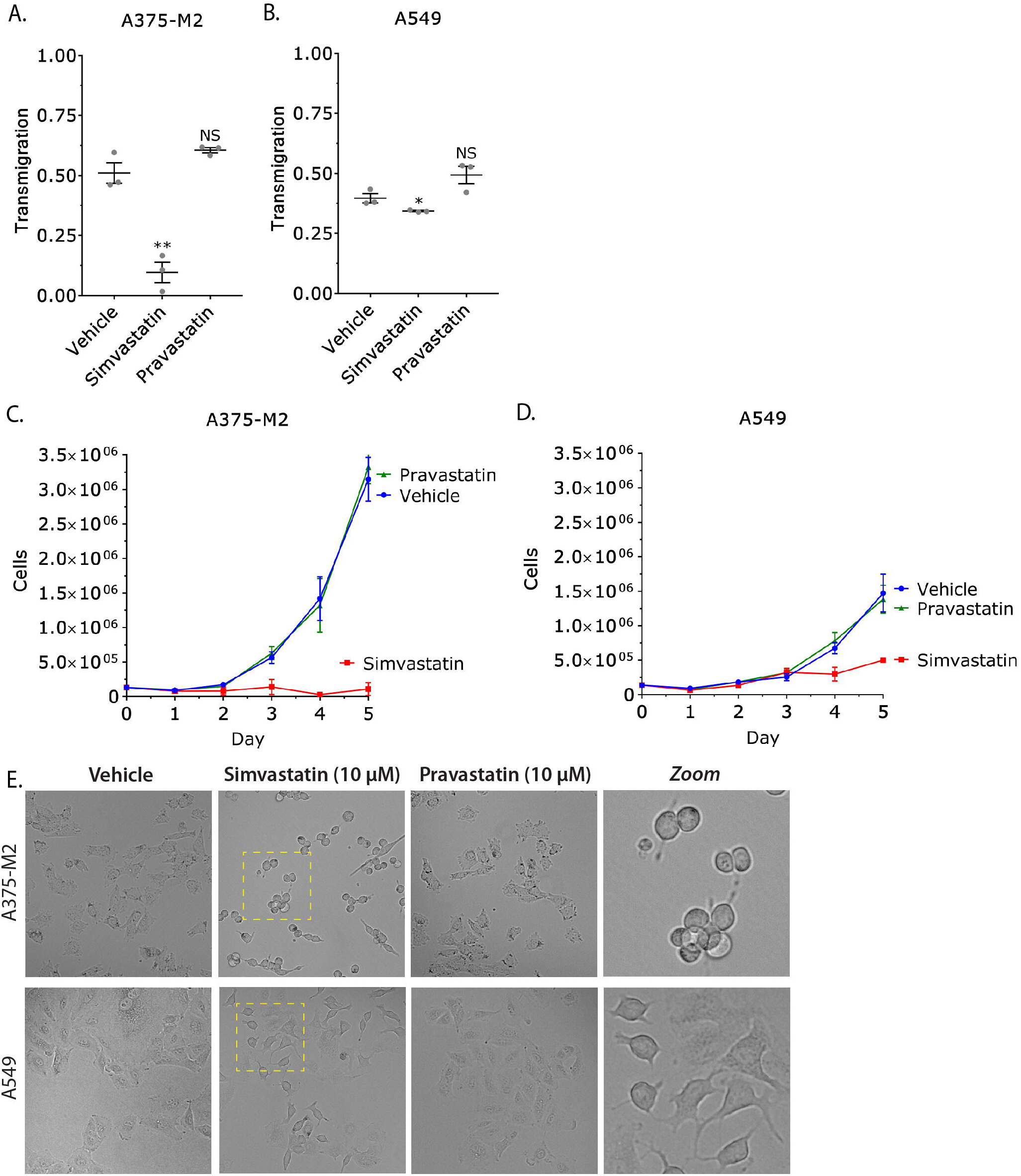
Simvastatin impairs the migration and proliferation of predominantly Vimentin expressing cells. **A-B.** Quantitative evaluation of transmigration for drug treated A375-M2 (A) and A549 (B) cells through fibronectin coated (10 µg/mL) polycarbonate filters with 8 µm pores. Statistical significance was determined by a two-tailed (unpaired) Student’s t-test. Error is SEM. **C-D.** Growth curves for drug treated A375-M2 (C) and A549 (D) cells on tissue culture plastic. Error is SEM. **E.** Brightfield images of A375-M2 (top) and A549 (bottom) cells one day after treatment with Vehicle, Simvastatin, or Pravastatin. All experiments were performed at least three times. * - p ≤ 0.05, ** - p ≤ 0.01, and *** - p ≤ 0.001

In order to evaluate if Simvastatin has a general effect on cancer cell motility, we subjected drug treated A375-M2 and lung cancer A549 cells, which also express Keratin (Fig. 1B), to transmigration assays. Using this approach, we found that Simvastatin decreases transmigration for A375-M2 (∼20% of control) and to a lesser extent A549 cells (∼85% of control), whereas Pravastatin did not have a significant effect (Fig. 4A-B). Because we observed a ∼50% increase in the number of apoptotic cells after Simvastatin treatment (Fig. S2C), we next determined if Simvastatin had a general effect on cell proliferation. To accomplish this, we counted cells over 5 consecutive days in order to generate growth curves for A375-M2 and A549 cells. Strikingly, we found that Simvastatin but not Pravastatin treatment inhibited the proliferation of both cell types (Fig. 5C-D). However, this effect was significantly more pronounced for A375-M2 cells, which predominantly express Vimentin (Fig. 1B). Consistent with Vimentin increasing the size and strength of focal adhesions, A375-M2 and A549 cells were frequently de-adhered one day after Simvastatin treatment (Fig. 5E) (1). Again, this effect was much more pronounced for A375-M2 cells. In contrast, cells treated with Pravastatin were not significantly different from Vehicle treated (Fig. 5E). Altogether, these results demonstrate that the migration and proliferation of cancer cells is inhibited by Vimentin bundling.

## Discussion

Here, we describe for the first time the contribution of the Vimentin intermediate filament network to LBBM. In contrast to mesenchymal migration, we demonstrate that depleting cells of Vimentin increases the speed of LBBM. A result that may be explained by our observation that Vimentin is entirely localized to the cell body, which is thought to resist the motile force produced by leader blebs. In agreement with this concept, leader blebs that have spontaneously separated from the cell body are extremely fast (3). Our measurements found leader bleb area to be increased after Vimentin RNAi, whereas the cell body area was reduced. These results suggest that Vimentin limits the compressibility of the cell body by cortical actomyosin, therefore more cytoplasm can flow into leader blebs to increase their size. Consistent with previous reports, our measurements of stiffness found that Vimentin protects the cell against compression (24). Vimentin may be particularly important for protecting the nucleus from compressive force, as several adaptor proteins have been reported to connect it to the nuclear envelope (25-27). Accordingly, Vimentin has been recently reported to protect the nucleus from rupture and DNA damage in confinement (28,29). In line with these results, we find the nucleus in Vimentin RNAi cells to undergo dramatic shape changes during LBBM. Moreover, cells depleted of Vimentin were found to more frequently undergo apoptosis. In mesenchymal cells, the density of the Vimentin network has been reported to inversely correlate with the actin retrograde flow rate in lamellipodia (30,31). Similarly, we speculate that the localization of Vimentin in the cell body may be important to not restrict the cortical actomyosin flow in leader blebs. Therefore, our work identifies Vimentin to be a fundamental regulator of LBBM.

Our results suggest that Vimentin increases the stiffness of the cell body to negatively regulate LBBM. To test this model, we used the cholesterol lowering statin, Simvastatin, which was identified after a screen of 1,120 biochemically active molecules to specifically bind and induce the bundling of Vimentin filaments (12). Indeed, using high-resolution live imaging of melanoma cells, we observed Vimentin to progressively bundle after Simvastatin treatment and not with another statin, Pravastatin. Because tissue culture cells obtain cholesterol from serum, this effect is not likely due to the lowering of cholesterol. In line with our model, Vimentin bundles led to the cell body being cemented in place, inhibiting LBBM. Using transmigration assays, which involves the migration of cells through 8 µm pores, we determined that Vimentin bundling could generally inhibit migration. However, this effect appears to be more pronounced in cells that predominantly express Vimentin. Additionally, the proliferation of cells predominantly expressing Vimentin was dramatically inhibited by Simvastatin. This result may be due to the loss of adhesion we observed after Simvastatin but not Pravastatin treatment. Thus, the migration and proliferation of cancer cells requires a flexible Vimentin network.

Our results are significant because in large cohort studies, patients taking statins for cholesterol have decreased cancer associated morbidity (22). However, statins are selectively localized to the liver, which is the major site of cholesterol production; therefore, this effect is more likely to be due to a decrease in blood cholesterol. In support of this idea, rapidly growing cancer cells require high uptake of extracellular cholesterol (32,33). Therefore, patients are not likely to benefit much from the effect of Simvastatin on Vimentin. Together with the work of others, our studies support the derivatization of Simvastatin for generating an entirely Vimentin selective molecule. Notably, mice lacking Vimentin exhibit few defects, therefore the systemic administration of a Vimentin selective molecule may be well tolerated (34-36). Thus, the perturbation of Vimentin function in cancer cells represents an attractive strategy for the prevention of metastasis.

## SUPPLEMENTAL INFORMATION

Supplemental information includes 2 figures and 4 movies and can be found with this article online.

## METHODS

### Cell culture

A375-M2 (CRL-3223) and A549 (CCL-185) cells were obtained from the American Type Culture Collection (Manassas, VA) and maintained for up to 30 passages in DMEM supplemented with 10% FBS (cat no. 12106C; Sigma Aldrich, St. Louis, MO), GlutaMAX (Life Technologies, Carlsbad, CA), antibiotic-antimycotic (Life Technologies) and 20 mM Hepes pH 7.4. Cells were plated on 6-well glass bottom plates (Cellvis, Mountain View, CA) either directly or after coating with 10 µg/ml human plasma fibronectin (cat no. FC010; Millipore, Billerica, MA), as noted in the figure legend.

### Pharmacological treatments

Simvastatin (cat no.1965) and Pravastatin (cat no. 2318) were purchased from Tocris Bioscience (Bristol, UK). DMSO (Sigma Aldrich) was used to make 1000X stock solutions for a working concentration of 10 µM. Prior to imaging under confinement, plates with PDMS slabs were incubated overnight in media with DMSO, Simvastatin, or Pravastatin. The following day, this media was replaced with fresh complete media containing DMSO, Simvastatin, or Pravastatin.

### Plasmids

Vimentin-FusionRed and H2B-FusionRed were purchased from Evrogen (Russia). mEmerald-Vimentin-7 and F-tractin-EGFP were gifts from Michael Davidson (Addgene; 54299) and Dr. Dyche Mullins (Addgene; 58473), respectively. FusionRed-F-tractin has been previously described (2). 1 µg of plasmid was transfected using a Nucleofector 2b device (Kit V; Lonza, Basel, Switzerland).

### LNAs

Non-targeting (cat no. 4390844) and Vimentin (cat no. 4390824; s14798) LNAs were purchased from Life Technologies. 50 nM (final concentration) LNA was transfected into cells using RNAiMAX (Life Technologies) diluted in OptiMEM (Life Technologies).

### Microscopy

Live high-resolution imaging was performed using a General Electric (Boston, MA) DeltaVision Elite imaging system mounted on an Olympus (Japan) IX71 stand with a computerized stage, environment chamber (heat, CO2, and humidifier), ultrafast solid-state illumination with excitation/emission filter sets for DAPI, CFP, GFP, YFP, and Cy5, critical illumination, Olympus PlanApo N 60X/1.42 NA DIC (oil) objective, Photometrics (Tucson, AZ) CoolSNAP HQ2 camera, proprietary constrained iterative deconvolution, and vibration isolation table.

### Confinement

This protocol has been described in detail elsewhere (16). Briefly, PDMS (Dow Corning 184 SYLGARD) was purchased from Krayden (Westminster, CO). 2 mL was cured overnight at 37C in each well of a 6-well glass bottom plate (Cellvis). Using a biopsy punch (cat no. 504535; World Precision Instruments, Sarasota, FL), an 8 mm hole was cut and 3 mL of serum free media containing 1% BSA was added to each well and incubated overnight at 37 °C. After removing the serum free media containing 1% BSA, 200 µL of complete media containing trypsinized cells (250,000 to 1 million) and 2 µL of beads (3.11 µm; Bangs Laboratories, Fishers, IN) were then pipetted into the round opening. The vacuum created by briefly lifting one side of the hole with a 1 mL pipette tip was used to move cells and beads underneath the PDMS. Finally, 3 mL of complete media was added to each well and cells were recovered for ∼60 min before imaging.

### Leader bleb, cell body, and Vimentin area measurements

For leader bleb, cell body, and Vimentin areas, freshly confined cells were traced from high-resolution images with the free-hand circle tool in Fiji (https://fiji.sc/). From every other frame, the percent of cell body area for leader blebs, percent of total for cell body areas, and percent of cell body area for Vimentin was calculated in Microsoft Excel (Redmond, WA). Frame-by-frame measurements were then used to generate an average for each cell. All statistical analyses were performed in GraphPad Prism (La Jolla, CA).

### Cell migration

To perform cell speed and directionality analyses, we used a Microsoft Excel plugin, DiPer, developed by Gorelik and colleagues and the Fiji plugin, MTrackJ, developed by Erik Meijering for manual tracking (37,38). For minimizing positional error, cells were tracked every other frame. Brightfield imaging was used to confirm that beads were not obstructing the path of a cell. All statistical analyses were performed in GraphPad Prism.

### Cell stiffness measurements

The gel sandwich assay has been described in detail elsewhere (3).

### Transmigration

Prior to transmigration assays, polycarbonate filters with 8 µM pores (cat no. 83-3428; Corning, Corning, NY) were coated with 10 µg/mL fibronectin (Millipore) by air drying for 1 hr. After permitting ∼100,000 cells in serum free media to attach (1 hr), DMSO, Simvastatin, or Pravastatin were added to each well. Bottom chambers contained 20% FBS in media to attract cells. After 24 hr, A375-M2 or A549 cells from the bottom of the filter were trypsinized and counted using an automated cell counter (TC20; Bio-Rad, Hercules, CA). Transmigration was then calculated as the ratio of cells on the bottom of the filter vs. the total. All statistical analyses were performed in GraphPad Prism.

### Growth curves

On day zero, ∼125,000 cells were plated in 6-well tissue culture plates in complete media with DMSO, Simvastatin, or Pravastatin. For 5 consecutive days, A375-M2 or A549 cells were trypsinized and counted using an automated cell counter (TC20; Bio-Rad). Each day, wells were supplemented with fresh media and drug till their day to be counted. All plots were generated using GraphPad Prism.

### Western blotting

Whole-cell lysates were prepared by scraping cells into ice cold RIPA buffer (50 mM Hepes pH 7.4,150 mM NaCl, 5 mM EDTA, 0.1% SDS, 0.5% deoxycholate and 1% Triton X-100) containing protease and phosphatase inhibitors (Roche, Switzerland). Before loading onto 4– 12% NuPAGE Bis-Tris gradient gels (Life Technologies), lysates were cleared by centrifugation. Following SDS-PAGE, proteins in gels were transferred to nitrocellulose membranes and subsequently immobilized by air drying overnight. After blocking in Tris-Buffered Saline containing 0.1% Tween 20 (TBS-T) and 1% BSA, primary antibodies against Vimentin, (cat no. 5741; Cell Signaling Technology, Danvers, MA), pan-Keratin (cat no. 4545; Cell Signaling Technology), IQGAP1 (cat no. 20648; Cell Signaling Technology), p-RLC (cat no. 3671; Cell Signaling Technology), or RLC (cat no. 8505; Cell Signaling Technology) were incubated with membranes overnight at 4C. Bands were then resolved with Horse Radish Peroxidase (HRP) conjugated secondary antibodies and a C-Digit imager (LI-COR Biosciences, Lincoln, NE).

### Statistics

All sample sizes were empirically determined based on saturation. Outliers were identified by the ROUT method in GraphPad Prism and excluded from further analyses. As noted in each figure legend, statistical significance was determined by either a two-tailed (unpaired) Student’s t-test, F test, or ordinary one-way ANOVA followed by a post-hoc multiple comparisons test. Normality was confirmed by a D’Agostino & Pearson test in GraphPad Prism. * - p ≤ 0.05, ** - p ≤ 0.01, and *** - p ≤ 0.001

## ACKNOWLEDGEMENTS

We thank members of the Logue Lab for insightful discussions and critical reading of this manuscript. We would also like to thank the administrative staff within the Department of Regenerative and Cancer Cell Biology at the Albany Medical College. This work was supported by start-up funds provided by the Albany Medical College.

## AUTHOR CONTRIBUTIONS

J.S.L. conceived and designed the study. S.M.T. performed all experiments except for the cell stiffness, Western blotting, transmigration, and growth curve assays after pharmacological treatments, which were performed by S.B.L. J.S.L. wrote the manuscript with comments from S.B.L. and S.M.T.

## COMPETING FINANCIAL INTERESTS

The authors declare no competing financial interests.

## SUPPLEMENTAL FIGURE LEGENDS

**Supplemental figure 1. Directionality for Vimentin RNAi and OE cells. A-B.** Quantitative evaluation of directionality over time for control (non-targeting) vs. Vimentin RNAi (A) and for EGFP alone vs. Vimentin-FusionRed (B) over-expressing A375-M2 cells under PDMS. Error is SEM.

**Supplemental figure 2. Survival for Vimentin RNAi and OE, and drug treated cells. A-C.** Quantitative evaluation of live A375-M2 cells after Vimentin RNAi (A) and OE (B), or after drug treatments (C) under PDMS.

**Supplemental movie 1.** Time-lapse imaging of an A375-M2 cell under PDMS transiently expressing Vimentin-mEmerald and H2B-FusionRed.

**Supplemental movie 2.** Time-lapse imaging of an A375-M2 cell under PDMS depleted of Vimentin by RNAi expressing EGFP-F-tractin and H2B-FusionRed.

**Supplemental movie 3.** Time-lapse imaging of an A375-M2 cell under PDMS treated with Vehicle (DMSO).

**Supplemental movie 4.** Time-lapse imaging of an A375-M2 cell under PDMS treated with Simvastatin (10 µM).

