## Supplementary figures and images for "A flexible network of Vimentin intermediate filaments promotes the migration of amoeboid cancer cells through confined environments"

### Supplemental figure 1

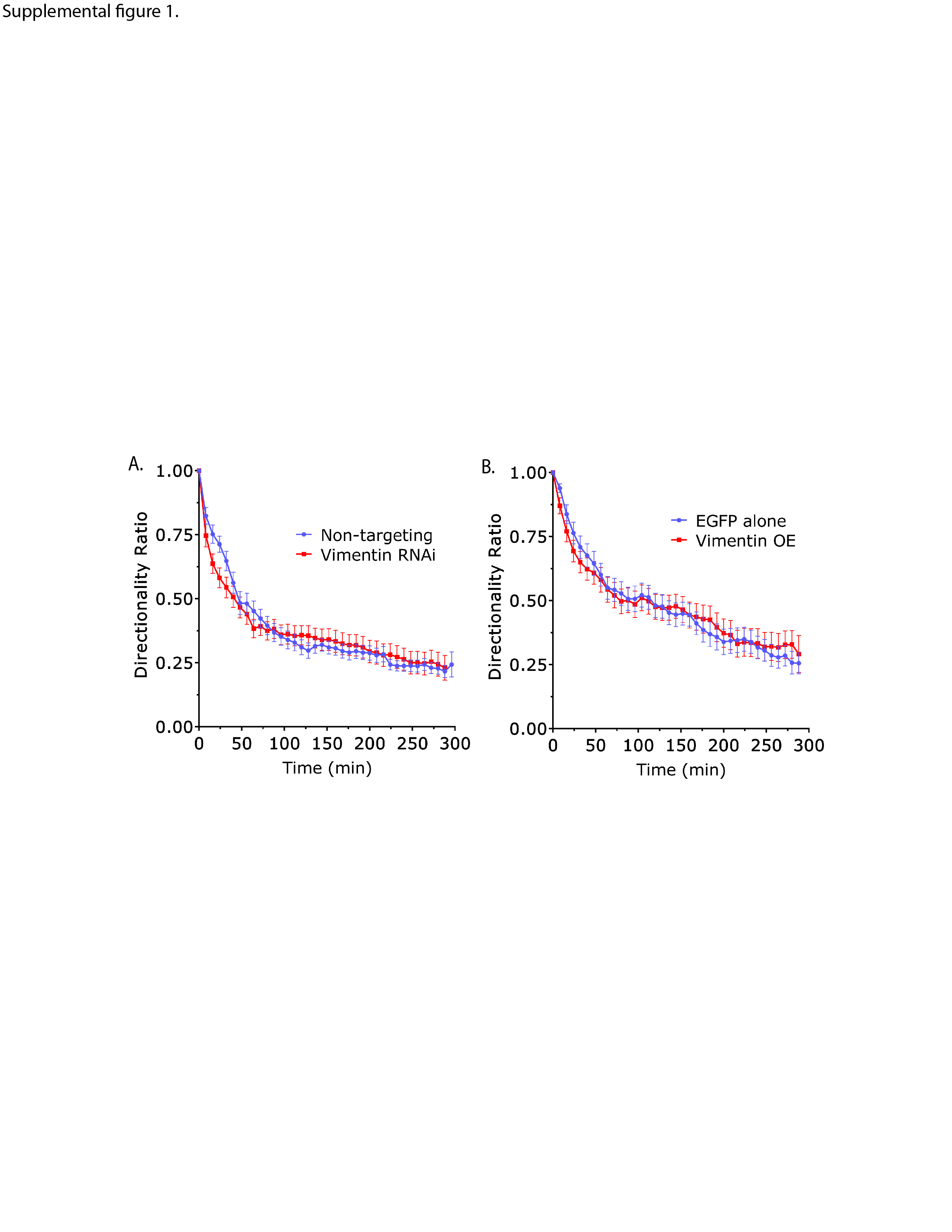

### Supplemental figure 2

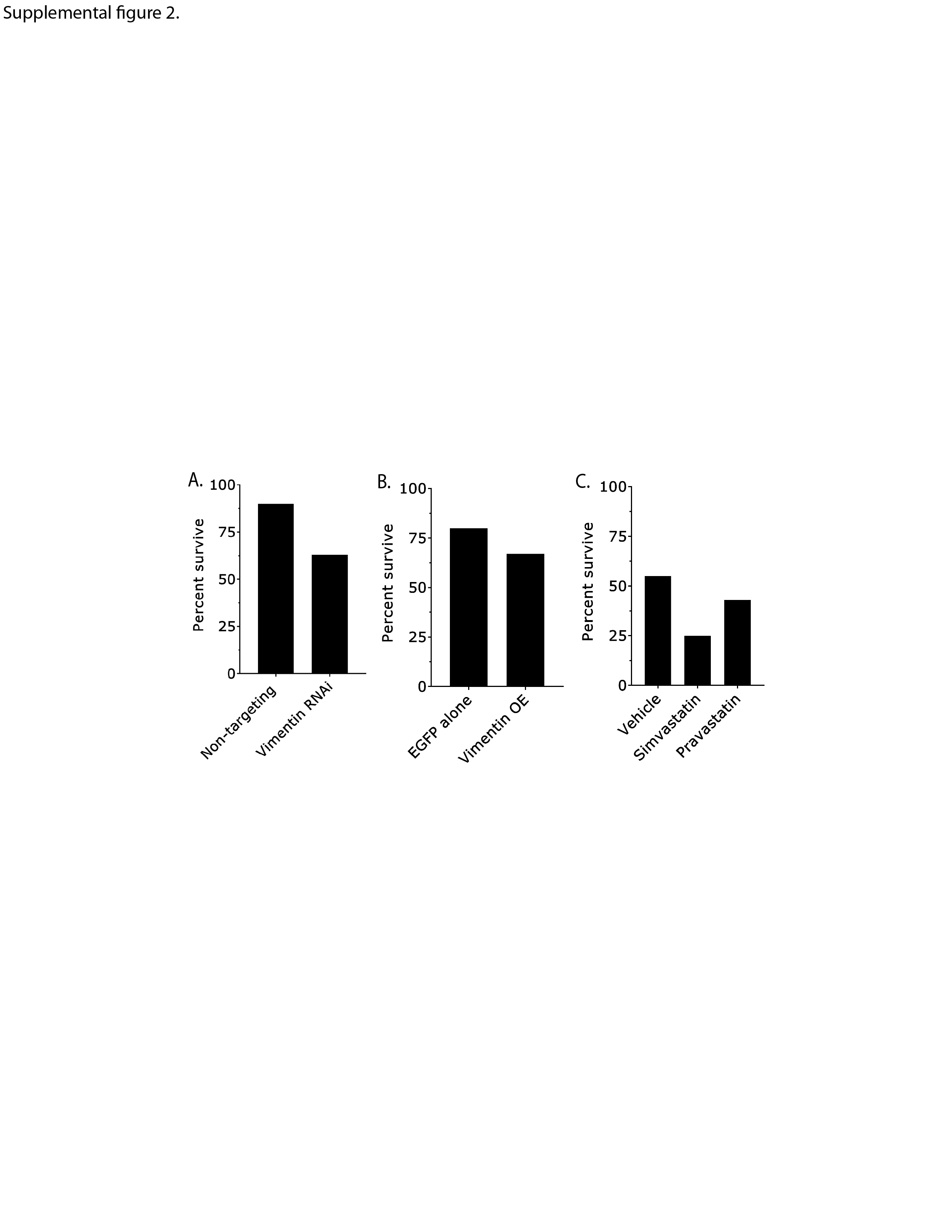
